# Non-redundant roles for paralogous proteins in the yeast glucose-sensing pathway

**DOI:** 10.1101/2025.10.07.680982

**Authors:** Yibo Si, Kshitiz Adhikari, Laura E. Herring, Daniel Isom, Shuang Li, Scott P. Lyons, Susan L. McRitchie, Blake R. Rushing, Susan J. Sumner, Henrik G. Dohlman

## Abstract

Paralogs engage in biological processes through both redundant and non-redundant functions. In the yeast *Saccharomyces cerevisiae*, approximately one-fifth of the genome consists of paralogs, with their encoded proteins involved in multiple pathways. However, the unique contributions of individual paralogs have remained poorly defined. Here, we undertook a systematic examination of eight paralog pairs in the glucose-sensing pathways, deleting each component and measuring the resulting changes in gene expression. To that end we established a new transcription reporter system to monitor the response to glucose as well as to non-preferred sugars in single cells. Focusing on the PKA catalytic subunits, comprised of the paralogs Tpk1 and Tpk3 as well as the isomorphic kinase Tpk2, we employed mass spectrometry to identify their contribution to cellular metabolism, used a GFP-based sensor to follow changes in cytosolic pH, and used BioID to identify unique and shared binding partners. Our data reveal that paralogs in the glucose-sensing pathway contribute in multiple and unique ways to signal transduction, and establish potential mechanisms driving the preservation of these and other duplicated genes throughout long periods of evolution.

## INTRODUCTION

Paralogs are pairs of genes that share the same parental gene [1], and they can be generated either through small-scale duplications or a whole genome duplication event [2–5]. Typically, one of the duplicated genes will eventually lose its function, due to an inactivating mutation or the loss of gene expression. Alternatively, both duplicated genes can be preserved, presumably because doing so provides some selective advantage for the organism. Paralogs that are functionally redundant can contribute to genetic robustness by maintaining proper growth in the face of genetic, regulatory or environmental perturbations [6–11]. For example, loss of a given paralog might lead to (or result from) a compensating change in the remaining paralog through alterations in gene transcription [12–15], subcellular localization [16–18], or protein stability and abundance [19, 20].

Paralogs can also evolve new functions that allow for adaptation to a broader set of environmental circumstances. Some paralogs are retained because they become complementary, wherein each inherits a subset of the functions of their parental gene (‘subfunctionalization’). Alternatively, each paralog can gain new functions that diverge from those of the original gene (‘neofunctionalization’) [21]. These alterations may be encoded genetically but can also arise through differences in post-translational regulation [22]. Our recent analysis revealed 3,500+ instances where only one of two paralogous proteins undergoes a post-translational modification, despite having retained the same target amino acid in both [23]. Thus, the preservation of a pair of paralogs can result from an interplay of multiple mechanisms that differ depending on their specific functions and the unique properties of the host organism [24–27].

A large number of paralogs emerged in the yeast *Saccharomyces cerevisiae* following a whole genome duplication event approximately 100 million years ago [28]; of the ∼6600 genes there are ∼550 paralog pairs annotated in the yeast genome [2, 3, 29]. The large number of paralogs in this organism provides a unique opportunity to study their contribution to genetic robustness, neofunctionalization and subfunctionalization. In addition, yeast is a single-cell organism that can exist stably in the haploid state [28]. Thus, it is possible to compare the functional consequences of deleting individual paralogous genes, in the absence of homologous genes and gene variants, in the same genetic and epigenetic background, and under identical environmental conditions.

Previous studies of specific paralog pairs in yeast have identified redundant or divergent functions in DNA replication, stress response and nutrient uptake. For example, some duplicated glycolytic genes have redundant functions that combine to contribute to high glycolytic flux; duplicated transcription factors cooperate to provide a stress response that is both stable and environmentally responsive within a cell population [30–32]. Moreover, a disproportionate number of these paralogs are components of the ribosome, or are involved in sugar sensing and sugar metabolism [29, 31, 33, 34]. Therefore, paralogs are likely to be particularly important in carbon source utilization leading to protein synthesis. Correspondingly, multiple paralogs are found in each of the two major glucose sensing pathways (Figure 1). One of the two pathways is initiated by the transporter-like receptor (transceptor) paralogs Rgt2 and Snf3 [35–37], Following glucose addition Rgt2 and Snf3 recruit the transcription repressor paralogs Mth1 and Std1 [38, 39], which are then phosphorylated by the casein kinase paralogs Yck1 and Yck2 [40]. These kinases also appear to phosphorylate Rgt2, but not Snf3, leading to stabilized expression [41]. Subsequent ubiquitination and degradation of these transcription factors leads to de-repression of low-affinity glucose transporters and the uptake of newly available sugars [38, 42–52]. Thus, the activity of this pathway is controlled by three pairs of paralogs: *RGT2*/*SNF3*, *YCK1*/*YCK2* and *MTH1*/*STD*1.

**Figure 1.**
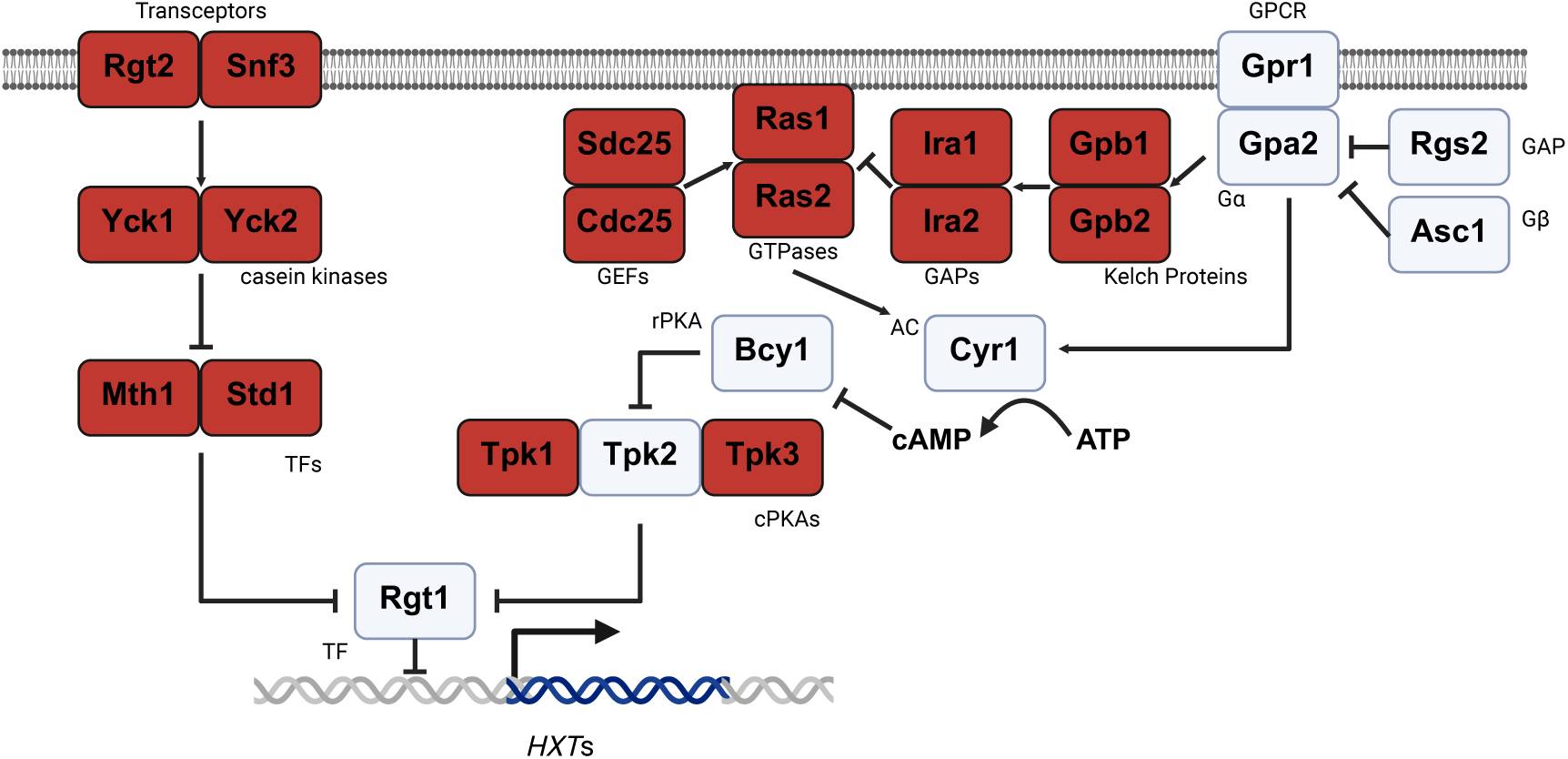
Yeast glucose sensing pathways. Two major pathways in *S. cerevisiae* sense extracellular glucose. All paralogous components are in dark red. GPCR: G-protein coupled receptor; GAP: GTPase-activating protein; GEF: guanine nucleotide exchange factor; AC: adenylyl cyclase; rPKA: regulatory subunit of protein kinase A; cPKA: catalytic subunit of protein kinase A; TF: transcription factor.

The second glucose sensing pathway is mediated by large and small G proteins, and is also enriched in paralogs. The large G protein is comprised of a typical Gα protein, Gpa2 [53], and an atypical Gβ subunit known as Asc1 (RACK1 in animals) [54]. Gpa2 is activated by the cell surface receptor Gpr1 [55–60], and inactivated by the GTPase accelerating protein Rgs2 [61].

The small G proteins, encoded by the paralogs *RAS1* and *RAS2* [62], are tuned by two pairs of paralogous proteins, the guanine nucleotide exchange factors Sdc25 and Cdc25 as well as the GTPase-activating proteins Ira1 and Ira2 [63–74]. Another pair of paralogous proteins, Gpb1 and Gpb2, regulate Ras1 and Ras2 through Ira1 and Ira2 [58, 75, 76]. Thus, the G protein-mediated signaling pathway has four paralog pairs: *SDC25*/*CDC25*, *RAS1*/*RAS2*, *IRA1*/*IRA2*, and *GPB1*/*GPB2*.

Finally, addition of glucose to starved cells upregulates the activity of Cyr1 (adenylyl cyclase) and results in a 2 to 3-fold spike of cAMP [53, 56, 75, 76]. Adenylyl cyclase is itself activated by Gpa2 as well as by Ras1 and Ras2 [62, 77–88], while Asc1 has the opposite effect [54]. cAMP then binds to the protein kinase A (PKA) regulatory subunit, Bcy1 [85], and triggers the release of catalytic subunits comprised of the paralogs Tpk1 and Tpk3, as well as the isomorphic kinase Tpk2 [89, 90]. These kinases go on to phosphorylate a panel of substrates involved in glucose uptake, metabolism and storage [89, 91–95], including Rgt1 leading to upregulation of low-affinity glucose transporters [44, 46, 47, 49, 50]. PKA-mediated phosphorylation can also regulate the paralogous transcription factors Msn2 and Msn4, the activities of which are highly associated with stress response [96–98]. Since glucose withdrawal is a strong stress for yeast, cAMP-PKA signaling is important for sensing and responding to a sharp transition of glucose availability [49, 99], as reviewed in [100].

Given the large number of paralogs in the glucose sensing pathways, and given our limited knowledge of paralog function, we undertook a systematic examination of each paralog pair in sugar sensing. Our approach was to delete individual paralogous components and measure changes in gene expression, metabolism and protein-protein interactions. As part of this effort, we established a new transcription reporter system to monitor the response to glucose as well as to non-preferred sugars. Focusing on the PKA catalytic subunits, we employed mass spectrometry to identify global metabolic changes and used a GFP-based sensor to follow metabolism-dependent changes in cytosolic pH. In this manner, we found multiple examples of non-redundant paralog pairs having differential effects in the response to glucose and other sugars. Finally, we used BioID (TurboID)-based proteomics to identify distinct binding partners of Tpk1, Tpk2 and Tpk3. These data reveal that paralogs in the glucose-sensing pathway contribute in unique ways to important biological processes, and establish potential mechanisms driving the preservation of these and other duplicated genes during the course of evolution.

## RESULTS

### Development of transcription reporters for yeast glucose sensing pathways

Many components of the two major glucose-sensing pathways in *Saccharomyces cerevisiae* are paralogs [29] (Figure 1). However, the evolutionary forces driving their retention, and their specific roles in glucose sensing, remain unclear. Paralogs may be maintained due to functional redundancy, which contributes to genetic robustness, or because they have diverged to perform non-redundant functions, through subfunctionalization or neofunctionalization [101]. However, while yeast glucose sensing is an attractive model for studying redundant and non-redundant functions of paralogs, a simple and quantitative measure of pathway activity has been lacking. Accordingly, we began by developing a transcription reporter system specific to the glucose-sensing pathways.

In our previous work, we compared the transcriptomes of wild-type cells after transient starvation and a 10 min treatment with either 0.05% glucose (‘low glucose’) or 2% glucose (‘high glucose’) [102, 103]. We were particularly interested in genes that are induced in low glucose or that are suppressed in high glucose, since we showed previously that glucose functions as an “inverse agonist” and prevents receptor-mediated cellular responses, and coordinates the programmed acidification of the yeast cytosol [104]. We cloned approximately 1 kb of the upstream regulatory region from each of 18 genes, all upregulated under low glucose conditions, and used these promoters to drive expression of the green fluorescent protein mNeonGreen. It is well established that intracellular pH is reduced under glucose-limiting conditions [104–107]; mNeonGreen, in comparison to GFP, is brighter, more stable, and less sensitive to pH [108]. The resulting reporter cassettes were fused with the *CYC1* terminator and integrated into a safe harbor locus on chromosome XVI [109]. Then, cells were grown in synthetic complete (SC) medium and 2% glucose to an OD_600_ of 0.8–1.2 and subsequently diluted to an OD_600_ of 0.02 in medium containing either 2% or 0.05% glucose. For each condition, cells were distributed into four wells of a half-area 96-well plate, and fluorescence intensity was measured every hour for 4 h using a single-cell imaging system. This approach allowed us to measure signaling intensity without the need to correct for cell density, while also providing information about the distribution of the response in the population. Of the 18 reporters tested, those with promoters for *HXT5* and *HXT7* consistently yielded high mNeonGreen expression in low glucose conditions.

Specifically, *pHXT5*- and *pHXT7*-driven mNeonGreen reporters showed 4-fold and 5-fold induction, respectively, following a shift to 0.05% glucose, and remained fully repressed in high glucose (Figure 2A; Supplemental Figure S1A). Thus, both reporters enable us to quantify the cellular response to glucose.

**Figure 2.**
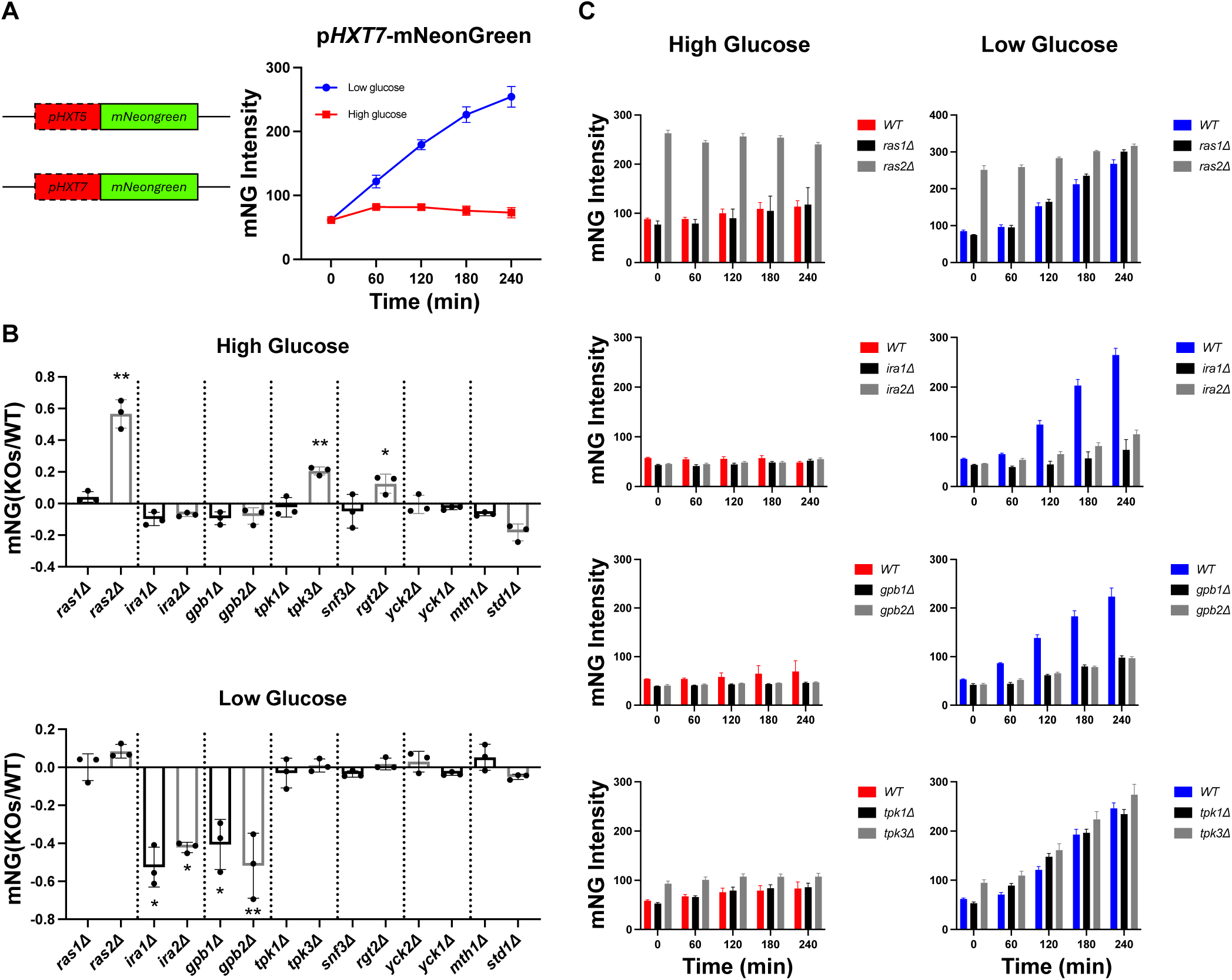
Paralogs have distinct roles in glucose sensing. **(A)** Left: promoters of *HXT5* and *HXT7* were selected to drive expression of mNeonGreen. The reporter cassette was integrated at *YPRCtau3* locus in Chromosome XVI. Right: Induction of *pHXT7*-mNeonGreen in wildtype BY4741 over 4 h. Low glucose: SC medium with 0.05% glucose; High glucose: SC medium with 2% glucose. **(B)** Transcription reporter assays using *pHXT7*-mNeonGreen. Wildtype and mutant strains were incubated in high or low glucose for 4 h. The reporter intensities from mutant strains were normalized to that from wildtype and transformed by log_10_. Error bars show standard deviation for three independent experiments; each symbol represents the mean of four wells. Two-tailed Student t-tests were done to compare intensities between mutants and wildtype. *: *p* < 0.05; **: *p* < 0.01. **(C)** Time course data for one representative experiment for deletions of *RAS1/2*, *IRA1/2*, *GPB1/2* and *TPK1/3*. Error bars show standard deviation for four wells representing the means of ∼1500 cells per well.

### Paralogs have distinct roles in glucose sensing

Having established transcription reporters for glucose sensing, we next sought to determine if the paralogs have redundant or non-redundant roles in this pathway. We deleted 14 genes (7 paralog pairs) from the Gpr1 and Rgt2/Snf3 pathways and monitored reporter fluorescence over a 4 h period, under low and high glucose conditions. We calculated the mean fluorescence intensity for each well and compared the results from paralog knockout strains with those of the wildtype control (Figure 2). Under high glucose conditions, the *pHXT7*-mNeonGreen signal increased appreciably in the *ras2*Δ*, tpk3*Δ and *rgt2*Δ strains. In contrast, deletion of the corresponding paralogs had little or no effect on reporter activity (Figure 2B&C). In low glucose conditions we observed a distinct pattern of activity; the absence of *IRA1*, *IRA2*, *GPB1*, and *GPB2* markedly impaired the induction of *pHXT7*-mNeonGreen. Loss of *YCK1*, *YCK2, MTH1, and STD1* had no effect in either condition (Figure 2B&C). Under both conditions, the *ras2*Δ strain exhibited constitutive induction, in keeping with its established role in coordinating and integrating the response to both receptor systems [102, 103]. In every case we obtained similar results for the *pHXT5*- and *pHXT7*-mNeonGreen reporters (Supplemental Figure S1). Therefore, by this measure paralogs in Gpr1/Ras signaling pathway are not redundant, while paralogs in Rgt2/Snf3 pathway are mostly redundant.

Whereas most deletions of pathway components are viable, at least one of the components (*CDC25*) is essential and cannot be deleted. As an alternative approach, we turned to the auxin-inducible degron (AID) system [110, 111]. Starting with the *pHXT7*-mNeonGreen reporter strain, we introduced the *osTIR1* gene and each paralog fused to AID. *osTIR*1 encodes an F-box protein, which is a part of the SCF ubiquitin ligase. When the plant hormone auxin is added it binds to OsTIR1, causing the SCF complex to recognize and recruit the AID tag on the target protein [112, 113]. This interaction then leads to ubiquitination of the tagged protein, marking it for rapid degradation by the proteasome.

Cells expressing the paralog-AID fusions were treated for 30 min with either 150 µM of the auxin 3-indoleacetic acid (IAA) or with DMSO. The cells were centrifuged and then resuspended in low or high glucose, with or without IAA, and reporter fluorescence was monitored for 4 h. Because of the difficulty of growing Ras1/Ras2-AID strains, we did not test them here. For each paralog-AID strain, we compared the fluorescence intensity in the presence and absence of IAA. Knock-down of Cdc25 resulted in a higher induction of the reporter, in both low and high glucose. In contrast, loss of Ira2 impaired the response in low glucose conditions, while knock-down of Ira1 had minimal effect (Figure 3). Knock-down of other paralogs had no effect, in either high or low glucose. Thus, use of the AID system reveals the importance of Cdc25, particularly in comparison to Sdc25. On the other hand, the knock-down of several paralogs (Gpb1, Gpb2, Tpk3, and Rgt2) failed to show differences even when they were evident following deletion of the genes. Taken together, our data show that at least six of eight paralog pairs are non-redundant in the glucose sensing pathway.

**Figure 3.**
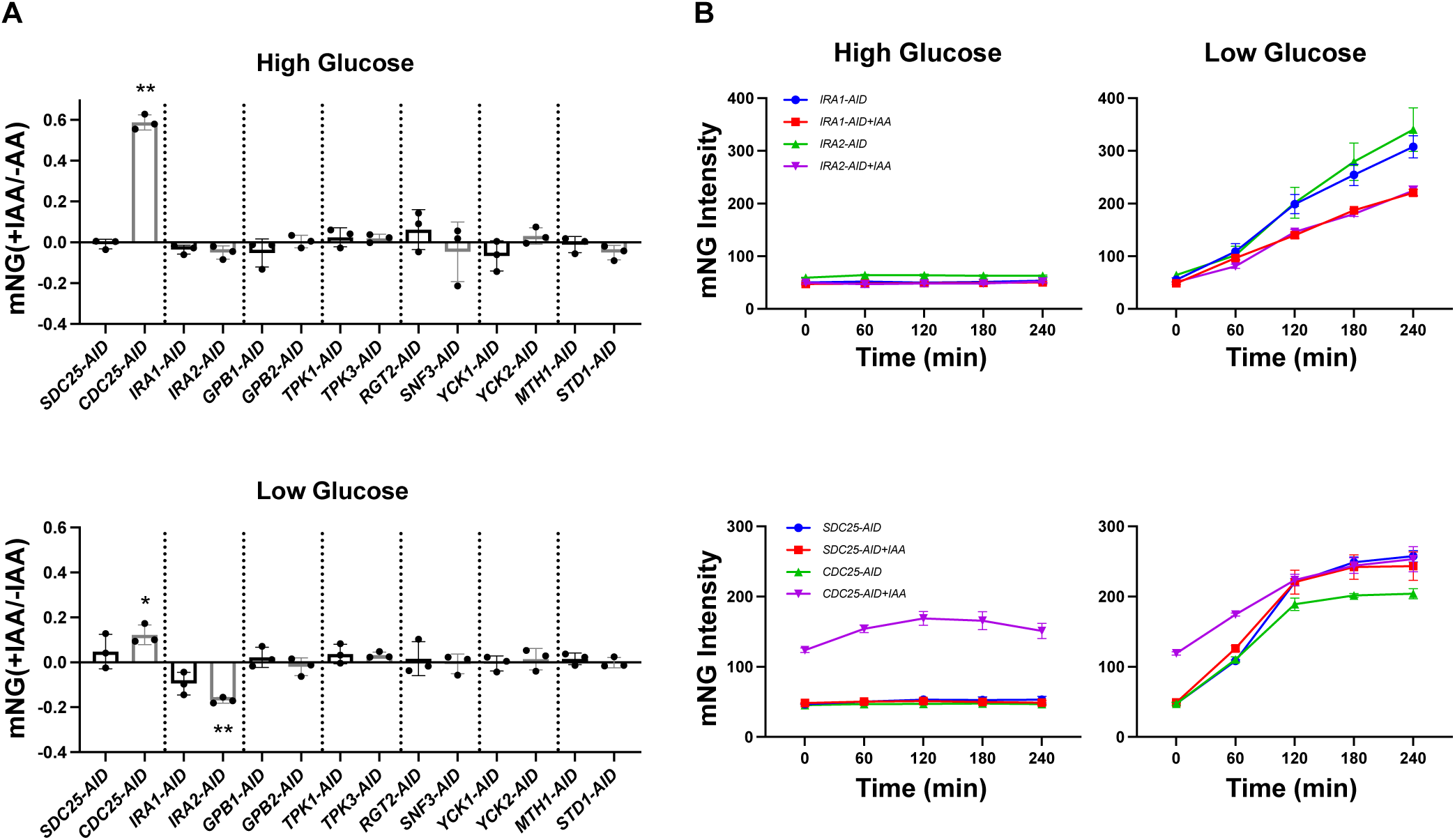
Protein degradation reveals unique roles of essential paralogs. **(A)** Transcription reporter assays using *pHXT7*-mNeonGreen. Paralog-AID strains were incubated in high or low glucose for 4 h, in the presence or absence of 150 μM IAA. The reporter intensities from each strain with IAA were normalized to that of the no IAA condition and transformed by log_10_. Error bars show standard deviation for three independent experiments; each symbol represents the mean of four wells. Two-tailed Student t-tests were done to compare intensities between paralog-AID fusion strains with and without IAA. *: *p* < 0.05; **: *p* < 0.01. **(B)** Time course data for one representative experiment for the transcription reporter assay in paralog-AID strains. Error bars show standard deviation for four wells representing the means of ∼1500 cells per well.

### Paralogs have distinct roles in signaling by non-preferred sugars

While glucose is a preferred carbon source, many other sugars can be consumed by *S. cerevisiae,* and doing so requires many of the same pathway components [60, 83, 104]. We next investigated whether paralogs in the pathway contribute differentially to the sensing of these non-preferred sugars (Figure 4). To that end, we measured the response to maltose, mannose, fructose, galactose and sucrose, all using the *pHXT7*-mNeongreen reporter (Figure 4A). Cells were grown in high glucose, washed and resuspended in SC medium containing one of the non-preferred sugars at 2%. Wild-type cells responded to maltose, galactose, and sucrose in a manner resembling that of low glucose, while mannose and fructose elicited weaker induction, resembling that of high glucose (Figure 4B). Depending on the stimulus conditions, we observed a distinct pattern of redundancy and non-redundancy when comparing individual paralog deletions. For example, whereas wildtype cells exhibited a time-dependent induction in response to maltose, galactose, and sucrose, deletion of *GPB1* or *GPB2* impaired the response to sucrose, but not to maltose or galactose (Figure 4A&B). We also identified non-redundant roles for paralogs that had previously appeared redundant. Deletion of *SNF3* did not alter the response to either high or low glucose, but reduced signaling in maltose and enhanced it in mannose and fructose (Figure 4A&C). Deletion of *TPK3*, but not *TPK1*, increased signaling in high glucose, while deletions of either *TPK1* or *TPK3* led to enhanced signaling in galactose and fructose (Figure 4A&B). Thus, paralogs that appeared redundant in their response to glucose had non-redundant functions in their response to non-preferred sugars. The differences between *TPK1* and *TPK3* were particularly notable because they were evident for a variety of sugars.

**Figure 4.**
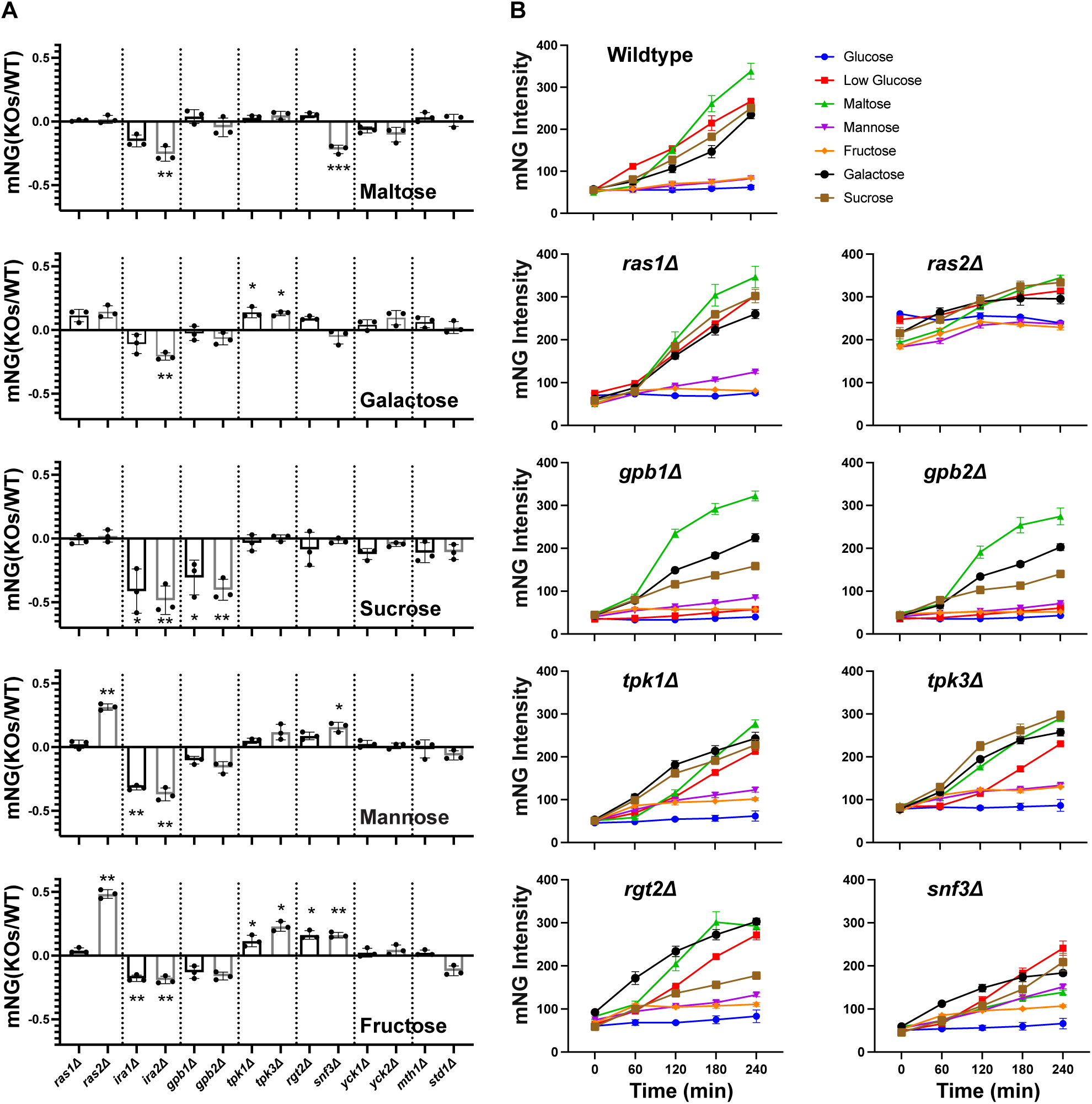
Paralogs have distinct roles in signaling by non-preferred sugars. **(A)** Transcription reporter assays using *pHXT7*-mNeonGreen. Wildtype and mutant strains were incubated in 2% of the non-preferred sugars (maltose, mannose, fructose, galactose or sucrose) for 4 h. The reporter intensities from mutant strains were normalized to that from wildtype and transformed by log_10_. Error bars show standard deviation for three independent experiments; each symbol represents the mean of four wells. Two-tailed Student t-tests were done to compare intensities between mutants and wildtype. *: *p* < 0.05; **: *p* < 0.01; ***: *p* < 0.001. **(B)** Time course data for one representative experiment for deletion of *RAS1/2*, *GPB1/2*, *TPK1/3* and *RGT2/SNF3*. Error bars show standard deviation for four wells representing the means of ∼1500 cells per well.

### Paralogous protein kinases have distinct roles in pH recovery

Glucose limitation causes a dramatic drop in intracellular pH [104–107]. In our previous work, we demonstrated that intracellular pH is regulated by multiple pathway components including the catalytic subunits of PKA, comprised of the paralogs Tpk1 and Tpk3 as well as the isomorphic kinase Tpk2 [104]. In the previous section we examined how these kinases regulate the response to different sugars. In this section, we consider how they contribute to the regulation of intracellular pH. To that end we used pHluorin, a pH-sensitive green fluorescent protein that provides a ratiometric measurement of intracellular pH in living cells [106, 114].

Each strain was transformed with the pHluorin plasmid, grown in 2% glucose, then washed in glucose-free medium and resuspended in buffered-medium containing either high or low glucose. Immediately after resuspension, intracellular pH was measured every minute for 25 min (Figure 5A). Under high glucose conditions, all *TPK* single-deletion strains exhibited pH recovery trajectories that were distinct from wildtype and from one another. Whereas all three of the mutant strains reached a final intracellular pH of approximately 0.2 units higher than that of wildtype, the *tpk2*Δ strain alone showed a delayed recovery. Under low glucose conditions, the *tpk1*Δ strain resembled wildtype. In contrast, *tpk3*Δ showed elevated intracellular pH while *tpk2*Δ exhibited decreased intracellular pH.

**Figure 5.**
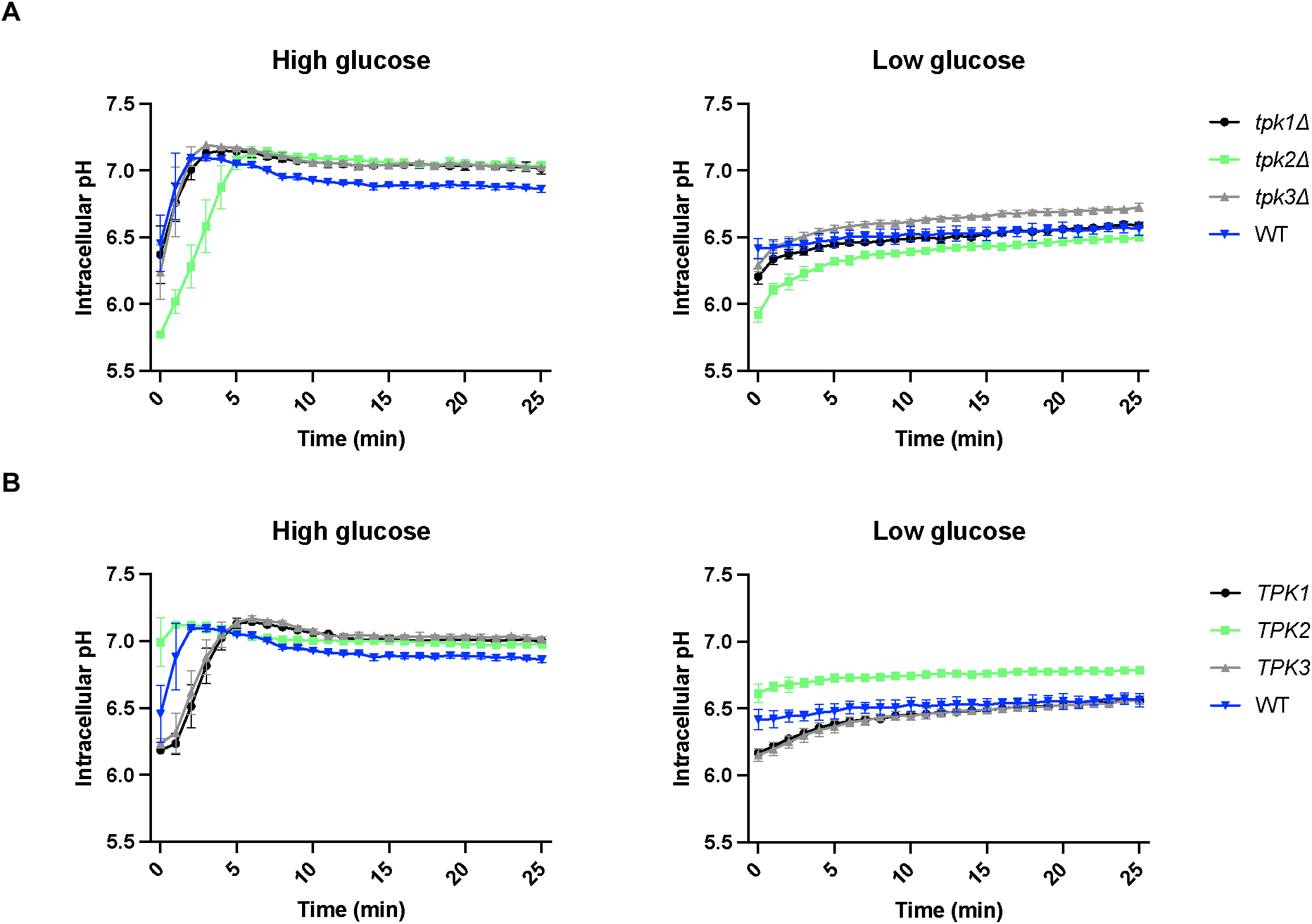
Paralogous protein kinases have distinct roles in pH recovery. **(A)** Kinetics of intracellular pH recovery in *TPK* single-deletion and wildtype strains. Left: High glucose conditions. Two-way ANOVA was performed to assess the effects of genotype (each *TPK* deletion vs. wildtype) on intracellular pH. All *TPK* deletions showed significant differences (interaction between genotype/time, *p* < 0.0001) compared to wildtype. Right: Low glucose conditions. *tpk2*Δ and *tpk3*Δ showed significant differences (*p* < 0.0001 for *tpk2*Δ, *p* < 0.0005 for *tpk3*Δ) compared to wildtype. **(B)** Kinetics of intracellular pH recovery in *TPK* double-deletion and wildtype strains. Left: High glucose conditions. All three *TPK*-only strains showed significant differences (*p* < 0.0001) compared to wildtype. Right: low glucose conditions. *TPK2-*only showed significant differences (*p* < 0.0001) compared to wildtype. For **(A)** and **(B)**, results for 26 time points (0 min to 25 min, 1 min interval) are plotted. Error bars show standard deviation for three independent experiments.

To further establish the individual contributions of each PKA, we examined pH recovery in double-deletion strains: *tpk2*Δ *tpk3*Δ (Tpk1-only), *tpk1*Δ *tpk3*Δ (Tpk2-only), and *tpk1*Δ *tpk2*Δ (Tpk3-only) (Figure 5B). In both high and low glucose, the Tpk1-only and Tpk3-only strains showed delayed recoveries. In low glucose, either Tpk1 or Tpk3 was sufficient to restore pH to wildtype levels, expression of Tpk2 alone resulted in pH levels approximately 0.2 units higher than that of wildtype. Taken together, these results indicate that the three Tpk proteins have different roles in responding to changes in glucose. Whereas Tpk2 promotes rapid recovery following glucose starvation, Tpk3 is responsible for maintaining proper pH over time.

### Paralogous protein kinases have distinct roles in metabolism

Our transcription reporter assays indicate that most paralogs in the Gpr1 pathway, most notably Tpk1 and Tpk3, have a distinct role in responding to glucose withdrawal (Figure 2). In the section above we show that the three Tpk isoforms also have distinct and important roles in reversing cellular acidification following a loss in glucose availability. The resulting decrease in ATP production leads to inactivation of the ATP-driven proton pump, Pma1, and a rapid increase in intracellular proton concentration [115–117]. Given that glucose, ATP, and pH play critical and multi-faceted roles in cellular metabolism [118, 119], we next asked how the kinases affect the abundance of a broader set of cellular metabolites.

For these experiments the three double deletion strains were placed in either high or low glucose for 2 min, and then collected for untargeted metabolomics via UHPLC high resolution mass spectrometry analysis as previously described [102, 103]. To assess metabolic changes, we compared each mutant strain to the wild type and calculated the fold change (FC) in metabolite abundance (with negative FC indicating wildtype > mutant). Differentially regulated metabolites (DRMs) were defined as those with an absolute FC > 2 and a *p*-value < 0.05 (Figure 6, Supplemental Dataset 1). Under both high and low glucose conditions, each mutant strain exhibited distinct DRMs. Venn diagrams illustrate the unique and shared DRMs across these three mutant strains (Figure 6A). None of the double deletion strains replicated the wildtype metabolic profile, which indicates that each single Tpk makes a unique contribution to glucose metabolism. The Tpk3-only strain showed the greatest number of DRMs in high glucose, whereas the Tpk2-only strain exhibited the greatest number in low glucose. These results indicate that Tpk3 has a comparatively minor role on metabolism under high glucose conditions while Tpk2 has a minor role under low glucose conditions.

**Figure 6.**
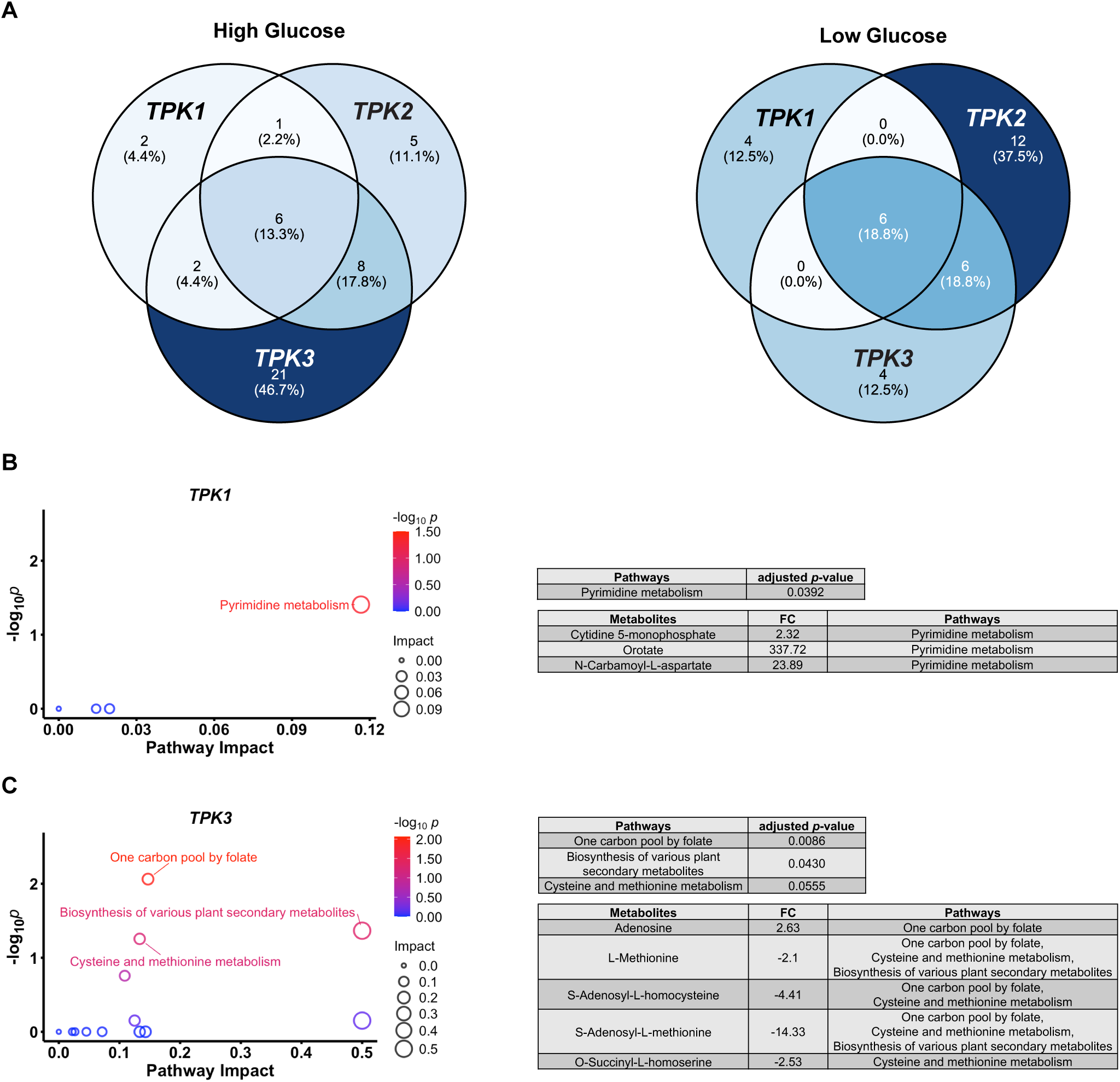
Paralogous protein kinases have distinct roles in metabolism. **(A)** Venn diagrams for differentially regulated metabolites (DRMs) in cells expressing Tpk1, Tpk2 or Tpk3 alone. Percentages: number of grouped DRMs / total DRMs × 100%. *TPK1*: Tpk1 only; *TPK2*: Tpk2 only; *TPK3*: Tpk3 only. High glucose: SC medium with 2% glucose; Low glucose: SC medium with 0.05% glucose. **(B)** and **(C)** Pathway analysis of enriched DRMs for *TPK1* and *TPK3* in high glucose condition. Left: scatter plots showing pathway impact calculated using MetaboAnalyst’s pathway topology analysis, which employs relative-betweenness centrality to assess the importance of matched DRM in each pathway. -log_10_*p*: adjusted *p*-values (false discovery rate calculated by Benjamini-Hochberg method) transformed by negative log_10_ for each pathway. Right: tables listing metabolites for significantly enriched pathways. FC: fold change calculated by comparing medians of all three *TPK*s and wildtype in six biological replicates (negative value means the abundance of DRM in wildtype is higher than that in *TPK*s).

To further investigate the unique role of each catalytic subunit in metabolism, we performed compound pathway analysis using MetaboAnalyst, a web-based platform for metabolomics studies (https://www.metaboanalyst.ca) [120]. In this application, we input all DRMs from each single-Tpk strain and queried for those over-represented in pathways annotated in the Kyoto Encyclopedia of Genes and Genomes (KEGG) [102, 103]. Under high-glucose conditions, pyrimidine metabolism was significantly affected in cells expressing only *TPK1*, with three metabolites (cytidine 5-monophosphate, orotate, N-carbamoyl-L-aspartate) in this pathway that were upregulated (Figure 6B; Supplemental Dataset 2). In cells expressing *TPK3*, we observed differences in five metabolites (adenosine, L-methionine, S-adenosyl-L-homocysteine, S-adenosylmethionine, O-succinyl-L-homoserine) involved in the one-carbon, cysteine, and methionine pathways (Figure 6C; Supplemental Dataset 2). The DRMs comparing wtildtype with Tpk1-only and with Tpk3-only were largely different (Figure 6B&C; Supplemental Dataset 1). Thus, Tpk1 and Tpk3 regulate distinct sets of metabolites and influence different metabolic pathways under high glucose conditions. No significantly enriched pathway (using the criteria of FDR < 0.05) was identified for the Tpk2-only cells in high glucose, or for any *TPK* strain in low-glucose conditions. These data indicate that Tpk2 is sufficient for proper metabolism under high glucose conditions and any Tpk isoform is sufficient under low glucose conditions. These observations indicate that Tpks have both shared and unique roles in metabolism. Moreover, differences are most evident when comparing the two paralogs, Tpk1 and Tpk3.

### Paralogous protein kinases have distinct binding partners

Given that Tpk1, Tpk2 and Tpk3 have similar catalytic activities [100], we hypothesized that the different cellular functions of Tpks are due to different protein interactions. To that end, we used a biotin-based proximity-labeling method to identify candidate interactors of each kinase subtype. By fusing a modified biotin ligase to the target protein, proteins associated with the fusion protein will be biotinylated upon biotin addition. The biotinylated proteins can then be identified by mass spectrometry [121–123]. We chose TurboID as the biotin ligase because of its high activity at 30°C, a suitable temperature for yeast culture [124–126]. We constructed strains by fusing TurboID to the C-terminus of each catalytic subunit, in an otherwise wildtype background. The cells were grown in SC + 2% glucose followed by the addition of 50 μM biotin. After 2 h, the cells were lysed and biotinylated proteins were enriched using magnetic streptavidin beads. The immobilized proteins were digested with trypsin, and peptides were identified and quantified by mass spectrometry. From the two independent experiments, we calculated the abundance of proteins enriched in the Tpk1-, Tpk2-, and Tpk3-TurboID strains, and compared each to that of wildtype. We considered proteins with log2 fold change > 1 and Student’s t-test *p*-value < 0.05 as significantly enriched (Supplemental Dataset 3).

As shown in Figure 7A, we identified 66, 120 and 101 proteins in the Tpk1-, Tpk2- and Tpk3-TurboID strains, respectively. Of the proteins identified, 35 (20.3%) were shared by all three Tpks; these include all three catalytic subunits as well as the PKA regulatory subunit, Bcy1 (Supplemental Dataset 3). The remaining interactors were unique to Tpk1 (9 proteins, 5.2%), Tpk2 (50 proteins, 29.1%) and Tpk3 (33 proteins, 19.2%). We then performed Gene Ontology (GO) term analysis for proteins enriched in each strain in g:Profiler [127], without including the PKA subunits themselves. Of all terms identified, half (107, 52.2%) were unique to Tpk2, while far fewer were unique to Tpk1 or Tpk3 or were shared by these two isoforms (Figure 7B). This is in line with the observation that Tpk2 has the most unique interactors. For the top 10 significantly enriched biological processes, distinct terms were identified for interactors of Tpk1 and Tpk2, while no biological process was determined for Tpk3 (Figure 7C). These results indicate that interactors of each Tpk are associated with unique cellular functions. Together, Tpk1, Tpk2, and Tpk3 exhibit distinct patterns of protein-protein interaction, and their interactors fall into different categories across Molecular Functions, Biological Processes, and Cellular Components (Supplemental Dataset 4).

**Figure 7.**
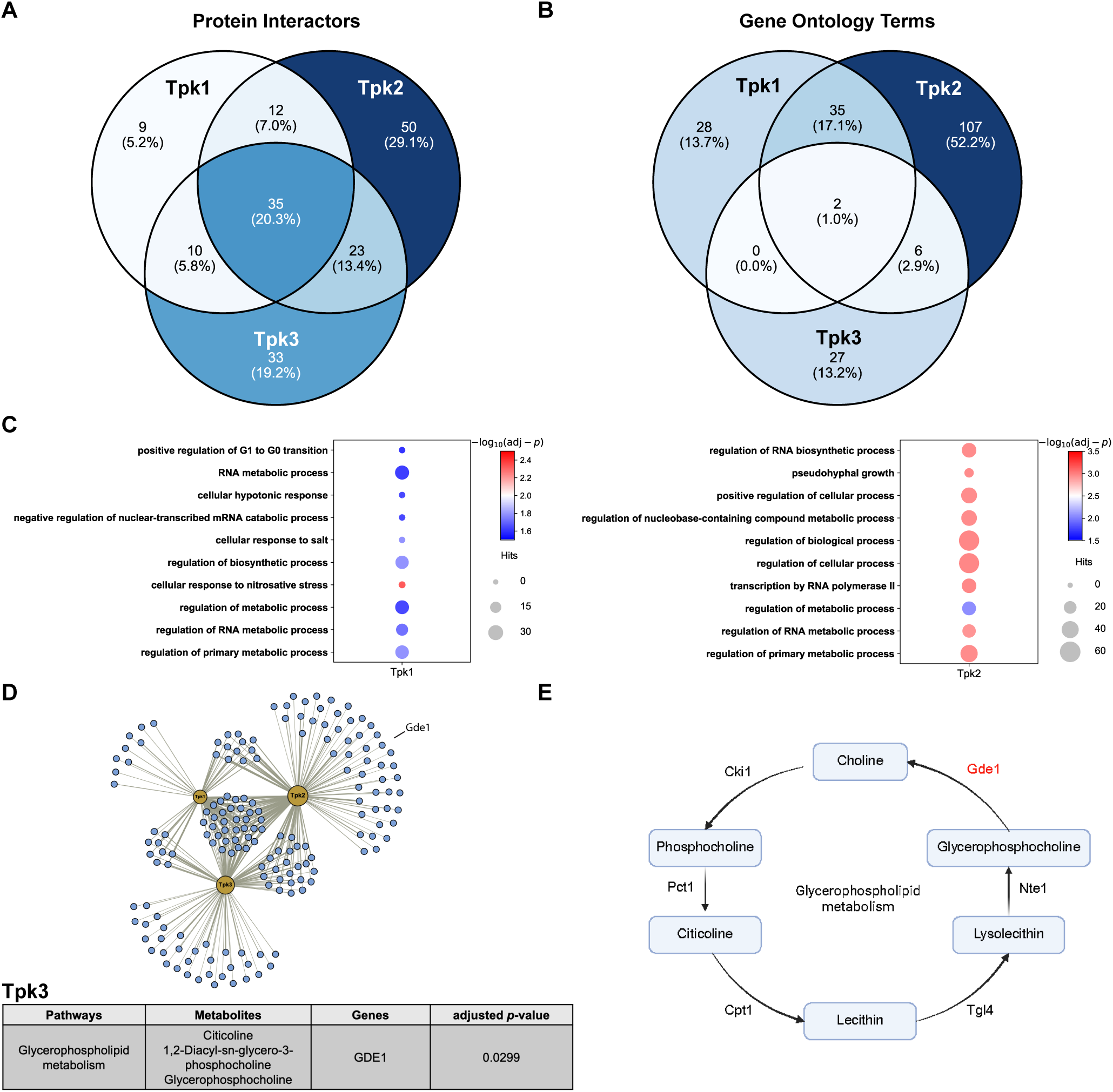
Paralogous protein kinases have distinct binding partners. **(A)** Venn diagram showing the number of proteins significantly enriched in the Tpk1- Tpk2- and Tpk3-TurboID strains. Percentages: number of grouped proteins / total proteins × 100%. **(B)** Venn diagrams showing the number of enriched GO terms for Tpk1- Tpk2- and Tpk3-interactors. Percentages: number of grouped GO terms / total terms × 100%. (C) Bubble plots showing the top 10 significant biological processes in Tpk1 and Tpk2, ranked by the adjusted *p*-values in each strain. **(D)** Top: protein-protein interaction network for Tpk1, Tpk2 and Tpk3. Yellow nodes represent Tpk1/2/3, blue circles represents binding partners. Sizes of nodes are scaled according to the number of interactors. Thicknesses of edges are scaled according to fold change in binding partner abundance compared to wildtype. Bottom: table listing a pathway enriched from integration analysis. **(E)** Simplified glycerophospholipid pathway and key enzymes are shown. Unique protein interactor highlighted in red.

To further explore the relationship between paralog functions in metabolism and in their protein-protein interactions, we performed integration analysis using the Joint Pathway Analysis module in MetaboAnalyst [102, 103]. By integrating data from metabolomics and proteomics analysis, we are able to identify functional relationships not evident from either analysis alone. We considered the differentially regulated metabolites (DRMs) and proteins associated with Tpk1, Tpk2, and Tpk3, and looked for instances where DRMs for one Tpk protein were functionally linked with interactors of the other Tpks. Our rationale was that any change in metabolite abundance observed in a single-Tpk strain result from the loss of protein interactions with the other two Tpks. To that end we grouped DRMs from Tpk1-only and unique protein interactors of Tpk2/3 and submitted them for joint pathway analysis. The abundance of three metabolites in the glycerophospholipid pathway were significantly reduced in the Tpk3-only strain (Figure 7D), while citicoline was downregulated in both Tpk1- and Tpk3-only strains (10.03-fold and 10.32-fold, respectively; Supplemental Datasets 1&5). Correspondingly, Gde1, which catalyzes the conversion of glycerophosphocholine to choline (Figure 7E) [128, 129], was identified as an interactor of Tpk2 but not Tpk1 or Tpk3 (Figure 7D). This suggests that the loss of interaction of Gde1 to Tpk2 leads to significant downregulation of citicoline in the Tpk1- and Tpk3-only strains. Together, Tpk1 and Tpk3 engage with distinct sets of protein interactors, and these distinct interaction patterns are strongly associated with unique metabolic profiles. This observation further supports the model that paralogs are non-redundant, and that functional differences arise from divergence in their protein-protein interaction networks.

Finally, we asked whether the identified protein interactors of Tpks are also phosphorylated, using a phosphoproteomic database of substrates for Tpk1/2/3, from Plank et al. [130]. In that study, phosphoproteomic analysis was performed to identify Tpk-dependent modifications at various time-points after the addition of 2% glucose to cultures growing in synthetic medium with glycerol as the carbon source. The Tpk isoforms were inactivated using low and high concentrations of kinase inhibitor. From a group of 121 candidate substrates, we identified 25 (20.6%) as significant interactors. These include Msn2, a core transcription factor involved in stress response [96–98], Net1, a core protein involved in cell cycle regulation [131–133] and Ume6, a transcription factor regulating meiosis, sporulation, carbon and nitrogen metabolism [134–136]. From 22 substrates that were transiently phosphorylated, six (27.2%) were included in our list of binding partners (Azf1, Ics2, Hbt1, Mbr1, Pfk26, Igo2; Supplemental Dataset 6). Of these, the 6-phosphofructo-2-kinase Pfk26 plays an important role in regulation of glycolysis [137–139]. Taken together, we conclude that Tpk1, Tpk2 and Tpk3 are not redundant in protein-protein interactions and cellular functions. Moreover, their distinct functions are tightly associated with their unique protein interaction profiles, and many of their interactors are substrates of phosphorylation.

## DISCUSSION

Duplicated genes are inherently unstable, and one or the other copy is likely to accumulate mutations and become a pseudogene or become eliminated entirely [24, 140, 141]. When they are retained, it is likely that the individual paralogs have acquired new functions, or else divided their duties to perform a subset of the functions of their common ancestor. The prevalence of duplicated genes in *Saccharomyces cerevisiae* has created an opportunity to investigate the evolutionary and selective pressures that favor paralog retention. Here, our results highlight the context-dependent nature of paralog function in glucose sensing and cell metabolism. Through a systematic analysis of individual gene deletion mutants we showed how each paralog pair contributes, in both shared and unique ways, to glucose sensing. By integrating new and well-established functional readouts, and by defining new binding partners for known pathway components, we have taken a major step by identifying distinct functions for seemingly redundant gene products.

Of the sixteen paralog pairs tested, all but *yck1/2* and *mth1/std1* altered transcription in response to a change in glucose availability. Under some conditions, deletion of one but not the other had an effect. Under other conditions, deletion of either paralog was sufficient. Moreover, the changes observed following gene deletion were not observed following induced protein degradation. These findings complement those from prior studies of other paralog pairs [12, 34, 142–148], or that relied on physiological changes such as growth and fermentation [10, 146, 149, 150]. Drawing clear conclusions from these past efforts has proven to be unexpectedly challenging, however. For example, a combined deletion of 13 single paralogs, each involved in the conversion of glucose to ethanol, revealed no defect with respect to gene expression, the formation of glycolytic products or growth in a variety of conditions [31]. Thus, establishing distinct functions for different paralogs can depend greatly on the pathway stimulus being applied and the functional output being measured.

Following our systematic analysis of paralogs in the glucose-sensing pathways, we turned our attention to the activities of the three catalytic subunits of PKA. Although any single *TPK* gene is sufficient to support cell growth [89], unequal roles have been noted in filamentous growth and mitochondrial biogenesis [91, 94, 95, 151, 152]. Other studies have considered differences in their regulation by phosphorylation [22, 69] and the resulting changes in catalytic activity [153] or subcellular localization [16–18] Our analysis reveals that the three catalytic subunits exhibit unique molecular interactions, and the distinct interactomes correspond to differences in substrate phosphorylation, as well as in their regulation of transcription and metabolism.

Our decision to investigate the glucose sensing pathways in yeast was based on a number of practical considerations. First, these pathways are enriched in paralogs. Moreover, the identities of pathway components, and their order of action, are well understood [154]. This allowed us to compare multiple individual paralogous proteins in their response to a single stimulus, and using a common transcriptional readout. By studying paralogs in the yeast glucose sensing pathways we have established important differences in their response to multiple stimuli, including non-preferred sugars. By integrating data from multiple approaches, including proteomics and metabolomics, we have gained a more thorough understanding of gene duplications that have persisted throughout long periods of evolution [102, 103]. More generally, our integrative approaches for investigating paralogs in the yeast glucose-sensing pathway could provide a guide for studies of paralogs involved in other processes, and may help to explain paralog retention in other organisms. For example, there are four PKA catalytic subunit isoforms in humans (PKACα, PKACβ, PKACγ and PRKX), and these have cell type-specific expression patterns that partially explain their retention [155]. However, the existence of three catalytic PKA subunits in a single cell organism like yeast implies that such isoforms have unique functions even within a given cell type. More broadly, the complex interplay between paralog pairs and their coordinated influence on cellular processes provides a strong rationale for the evolutionary retention of multiple subtypes of functionally similar proteins.

## MATERIALS & METHODS

### Plasmids & Primers

All plasmids used in this study are listed in Supplemental Table 1. Transcription reporters were constructed in pRS406 [156]. The backbones were amplified using primer sets ‘pRS406-linearization-F1/ pRS406-linearization-R1’ and ‘pRS406-linearization-F2/ pRS406-linearization-R2’; the coding sequence of mNeonGreen was amplified from pYEplac181-mNeonGreen using primer set ‘mNG-F/mNG-R’; two homology arms were amplified from BY4741 genomic DNA using primer sets ‘HA-down-F/ HA-down-R’ and ‘HA-up-F/ HA-up-R’. All fragments, except promoters, were linked by HiFi assembly (NEB, #E2621S). 1000 base pairs of the upstream promoter region of *HXT5* and *HXT7* were amplified from BY4741 genomic DNA using primer sets ‘pHXT5-F/pHXT5-R’ and ‘pHXT7-F/pHXT7-R’, containing EcoRI/XmaI restriction sites. Amplified promotors were digested by restriction enzymes (EcoRI/XmaI) and inserted upstream of mNeonGreen by ligation. Primers related to this study are listed in Supplemental Table 2.

### Yeast Strains & Media

Yeast strains used in this study are listed in Supplemental Table 3. Both transcription reporters, in tandem with an *URA3* selection marker, were amplified from *pHXT5*-mNeonGreen and *pHXT7*-mNeonGreen using primer set ‘HA-up-F/HA-down-R’. Amplified reporter cassettes were integrated in a safe harbor at chromosome 16 [109] in BY4741 by lithium acetate method [157]. All single gene deletions were created using PCR amplification of *KanMX6* from pFA6-KanMX6 with primer sets listed in Supplemental Table 2 followed by transformations and G418 selection [158]. Integrations of reporters and *KanMX6* were confirmed by PCR-based genotyping using primer sets listed in Supplemental Table 2. The *tpk* double mutants were constructed by replacing each target gene with KanMX6. The double mutants were constructed by switching the mating type of one strain from *MAT**a*** to *MAT*α, with HO expressed from a plasmid, and then mating to an isogenic *MAT**a*** strain containing the second mutation. The diploid was then sporulated and spore products with the double knock-outs were confirmed with PCR [102]. The AID strains were constructed as C-terminal fusions to IAA7, as described [159]. The AID fragment was amplified from pM66 using primer set ‘pM66-F2/pM66-R1’; all homology arms for paralogs were amplified from BY4741 genomic DNA using primer sets listed in Supplemental Table 3. AID fragments and homology arms were ligated by HiFi assembly, and ligation products were used for transformation. The TurboID strains were constructed as C-terminal fusions to biotin ligase, as described [126]. TurboID cassettes for *TPK1/2/3* were amplified from pFA6a-TurboID-3MYC-natMX6 (Addgene, #126050) using primer sets ‘Tpk1-TurboID-C-F/Tpk1-TurboID-C-R’, ‘Tpk2-TurboID-C-F/ Tpk2-TurboID-C-R’ and ‘Tpk3-TurboID-C-F/Tpk3-TurboID-C-R’. Following transformations, AID and TurboID strains were confirmed by PCR using primers listed in Supplemental Table 2 and western blots described in the next section.

Cells were grown in synthetic complete (SC) medium with 0.05% glucose, 2% glucose or 2% non-preferred sugar where specified. SC: 1.7 g/L yeast nitrogen base (BD #DF0335-15-9), 5 g/L ammonium sulfate (Sigma-Aldrich #A4418), 30 mg/L adenine hydrochloride hydrate (Sigma-Aldrich, #A8751), 1 pellet/L sodium hydroxide (Sigma-Aldrich #221465), and either 0.69 g/L CSM-LEU mixture (MP Biomedicals, #114510512-CF) or 0.77 g/L CSM-URA mixture (MP Biomedicals #1145112-CF) and supplemented with uracil (Sigma-Aldrich, #U0750) when needed. Solid media contain 15% agar. For intracellular pH measurement, buffered-SC medium was used: 50 mM dibasic potassium phosphate (Sigma-Aldrich, #P3786), 50 mM dibasic sodium succinate (Sigma-Aldrich #14160) and medium was titrated to pH 5 with HCl.

### Immunoblotting

Cells were cultured in SC + 2% glucose to OD_600_ = 1.0-1.2 before harvest. For AID strains, 7.5 μL 100 mM 3-Indoleacetic acid (Sigma-Aldrich, #I3750) or DMSO alone were added into 5 mL cell culture 30 min before harvest. 5 mL cell culture was mixed with 263 μL trichloroacetic acid (Sigma-Aldrich, #T0699) and collected by centrifugation (2100 g, 2 min, 4°C). Cells were resuspended with 200 μL TCA buffer (10mM Tris-HCl pH 8.0, 10% TCA, 25 mM NH_4_OAc, 1 mM Na_2_EDTA, freshly prepared). The resuspensions were centrifuged again at 16,000 g for 10 min at 4°C. The supernatants were removed, and pellets were resuspended with 50-100 μL resuspension buffer (100 mM Tris-HCl pH 11, 3% SDS), heated at 95°C for 10 min and centrifuged at 16,000 g for 1 min. 95 μL supernatant was collected, and protein concentrations were measured by DC protein assay (Bio-Rad, #500-0113). For each genotype, 15 μg protein sample was resolved by 7.5% SDS-PAGE and transferred at 4°C (25 V, 16 h) to nitrocellulose membranes. V5 epitope tag (for AID) was detected by V5 antibody (Invitrogen, #R960-25) at 1:5,000. Myc epitope tag (for TurboID) was detected by Myc antibody (Cell Signaling, #2276) at 1:5,000. G6PDH antibody (Sigma-Aldrich, #A9521) was used at 1:10,000 as a loading control.

Donkey anti-rabbit-HRP (Jackson ImmunoResearch, #711035152) and goat anti-mouse-HRP (Promega, #W4021) were used at 1:10,000 as secondary antibodies. All antibody dilutions were prepared in 5% non-fat milk in TBST (100 mM Tris-HCl pH 7.5, 150 mM NaCl, 0.1% Tween-20). Immunoreactive species were detected by Clarity Western ECL kit (Bio-Rad, #170-5061) and imaged by ChemiDoc MP System (Bio-Rad).

### Transcription Reporter Assays

All data for transcription reporter assays were collected by single-cell imaging on a Celigo imaging cytometer (Revvity). Reporter strains were grown in 5 mL SC medium with 2% glucose to an OD_600_ = 0.8-1.2. Cells were harvested by centrifugation (2,100 g, 3 min, room temperature) and washed with TE buffer (100 mM Tris-HCl pH 7.5, 10 mM EDTA) twice. Cells were resuspended in 5 mL either SC + 2% glucose (high glucose) or SC + 0.05% glucose (low glucose). Resuspensions were diluted to OD_600_ = 0.02 in reservoirs with SC medium supplemented with 0.05% glucose, 2% glucose or 2% non-preferred sugars. 50 μL of diluted cell samples were added into a 96-well half-area plate (Greiner Bio-one, #675090) with four wells for each condition.

For AID strains, cells were treated with 3-Indoleacetic acid (100 mM stock in DMSO) or DMSO for 30 min before harvest. Cells were washed as described above, and resuspended in SC + 2% glucose + 150 μM IAA or SC + 0.05% glucose + 150 μM IAA. All samples were diluted and measured as described above.

The fluorescence intensity of each single cell was measured under the green channel in the Celigo every hour up to 4 h. Single cells were identified by the instrument, and mean intensity was determined for each well. Mean intensities from 4 wells were used for statistical analysis. At least three independent experiments were done for each condition. Paired t-test was conducted to compare differences in fluorescence intensity between paralog deletion strains and wildtype.

### Intracellular pH measurement

Intracellular pH measurements using the pH biosensor pHluorin was described in [104]. Briefly, pYEplac181-pHluorin was transformed into BY4741 *TPK* single- or double-deletion strains (*tpk1Δ*, *tpk2Δ*, *tpk3Δ, tpk1Δ tpk2Δ, tpk1Δ tpk3Δ* or *tpk2Δ tpk3Δ*). Transformed cells were cultured in 5 mL SC with 2% glucose to OD_600_ = 1, centrifuged at 2,100 g for 3 min at room temperature, washed twice with buffered-SC medium without glucose. Washed cells were resuspended with buffered-SC media containing 1.6% or 0.05% glucose, and 200 μL cells were added into a 96-well plate. The plate was placed in a SpectraMax fluorescence plate reader to measure intracellular pH kinetics over 25 min (22°C). As described in [104], the emission ratio at 520 nm responding to excitation at 395 nm and 480 nm (ratio = emission^395^/emission^480^). The ratio was used to calculate the intracellular pH according to a standard curve. The intracellular pH recovery of each deletion strain was compared to that of wildtype by two-way ANOVA.

### Sample Preparation for Metabolomics

Collection of samples for metabolomics analysis was described in [102, 103]. Briefly, cells were cultured in SC + 2% glucose overnight to reach OD_600_ = 1. Cultures were centrifuged, washed with TE buffer, resuspended with 10 mL SC + 0.05% glucose, and incubated at 30°C with shaking for 1 h. Glucose was added to a final concentration of 2% or 0.05% 2 min before harvest. 3 mL cells were mixed with 45 mL cold methanol and put on dry ice for 5 min. Cells were centrifuged and stored at -80°C. The pellets were resuspended with 80% methanol and transferred to 2 mL screw-capped tubes prefilled with ceramic beads (Roche, # 03358941001). The resuspensions were then homogenized twice using Bead Mill 24 Homogenizer (Fisher Scientific, #15-340-163) at 6 m/s for 40 s in two cycles at room temperature. After homogenization, samples were centrifuged at 16,000 g for 10 min at 4°C, and 500 μL supernatant was transferred into low-bind 1.7 mL tubes. Quality control (QC) study pools were created by combining an additional 65 μL from each sample and then aliquoting into tubes at a volume of 500 μL each. Method blanks were created by adding 80% methanol to empty homogenization tubes and processing in an identical manner to the study samples. All samples, blanks, and QC pools were dried overnight by Speedvac vacuum concentrator (ThermoFisher Scientific) and resuspended with 100 μL reconstitution buffer (95% water, 5% methanol with 500 ng/mL tryptophan d-5). The resuspended extracts were then centrifuged at 16,000 g for 4 min at room temperature. Supernatants were transferred into autosampler vials for LC-MS where 5 μL was injected for analysis. The study samples were randomized and interspersed with the blanks and QC pools.

### Metabolomics Data Capture via UHPLC High Resolution Mass Spectrometry

LC-MS metabolomics analysis was conducted as described previously [102, 103]. Briefly, metabolomics data were collected using a Vanquish UHPLC coupled to a QExactive HF-X Hybrid Quadrupole-Orbitrap Mass Spectrometer (ThermoFisher Scientific). Progenesis QI (Waters Corporation) was used for data processing. Background signals were removed by filtering out signals that did not have an average intensity fold change ≥ 3 in the QC study pools compared to the blanks. Data were then normalized by Progenesis QI (version 2.1, Waters Corporation). Peaks were identified and annotated by matching to both an in-house library and to public databases. Compound annotation and pathway analysis was done by the MetaboAnalystR 6.0 in R (https://www.metaboanalyst.ca/docs/RTutorial.xhtml). Metabolomics data from *tpk* double mutants were normalized to that from wildtype strain in both high and low glucose conditions. For any compounds annotated, median intensity (n = 6) was compared to that in wildtype to generate a fold change (FC). Differentially regulated metabolites (DRMs) were defined as compounds of which |FC| ≥ 2 and Wilcoxon rank sum test *p*-value < 0.05. Metabolic pathway analysis was performed on MetaboAnalyst website (http://metaboanalyst.ca, module ‘Pathway Analysis’). DRMs from each strain were submitted as compounds with settings:

*Enrichment method = Hypergeometric Test, Topology measure = Relative-betweenness Centrality, Reference metabolome = Use all compounds in the selected pathway*.

### Streptavidin-Western Blotting

All TurboID and wildtype strains were cultured in 50 mL SC + 2% glucose overnight to reach OD_600_ ∼ 0.5. Cells were centrifuged and washed in TE buffer before resuspension in SC + 2% glucose. Biotin (Sigma, #B4501; prepared in DMSO to 100 mM) was added to a final concentration of 50 μM and cells were then incubated for additional 2 h at 30°C. Whole cell extractions were collected using a protocol modified from that described in [124]. After incubation, cell cultures were collected by centrifugation (2,100 g, 10 min, 4°C), washed twice by wash buffer (10 mM Tris-HCl pH 7.5, 50 mM NaCl, 1 mM EDTA, 0.5 mM phenylmethylsulfonyl fluoride), and pelleted (2,100 g, 10 min, 4°C). The pellets were resuspended with 600 μL lysis buffer (20 mM HEPES-OH pH 7.5, 10 mM MgCl_2_, 0.4 M NaCl, 2 mM EGTA, 10% glycerol) supplemented with protease inhibitor cocktail (ThermoFisher Scientific, #78340). The resuspensions were transferred to 2 mL screw-top microtubes (Revvity, #19-622), and cells were broken using a Bead Mill 24 Homogenizer at 6 m/s for 30s (4 cycles, 4°C). After homogenization, microtubes were centrifuged at 16,000 g for 10 min, and the supernatant was collected. 2 μL benzonase (Sigma-Aldrich, #70664) was added into the supernatant followed by 20 min incubation at 4°C. After incubation, tubes were centrifuged at 16,000 g for 10 min, and the supernatant was collected as protein samples. 15 μg protein from TurboID and wildtype strains was resolved by 7.5% SDS-PAGE and transferred at 4°C (25 V, 16 h) to nitrocellulose membrane. The membrane was blocked by 2% BSA in TBST for 1 h at room temperature and incubated with Ultra Streptavidin-HRP (ThermoFisher Scientific, #N504) at 1:20,000 in 2% BSA for 40 min at room temperature. The membrane was washed three times using TBST and biotinylated proteins were detected by chemiluminescence as described above.

### Sample Preparation for TurboID-based Proteomics

50 mL cultures from TurboID and wildtype strains were harvested and whole cell extractions were collected as described above. For each sample, 1.2 mg protein was diluted into 1 mL lysis buffer with protease inhibitor cocktail. 60 μL Pierce streptavidin magnetic beads (ThermoFisher Scientific, #88817) were added to capture biotinylated proteins and incubated overnight at 4°C. Beads were collected by DynaMag^TM^-2 Magnet (ThermoFisher Scientific, #12321D) and washed 10 min with 1.5 mL wash buffer 1 (lysis buffer + 0.1% Triton X-100 + protease inhibitor cocktail) twice by rotating tubes at 4°C. Beads were then washed once for 10 min with wash buffer 2 (20 mM HEPES-OH pH 7.5, 10 mM MgCl_2_, 1.5 M NaCl, 2 mM EGTA, 10% glycerol, 0.1% Triton X-100, 1 mM DTT) at 4°C, once for 10 min with wash buffer 3 (10 mM Tris-HCl pH 8, 2 M urea) at room temperature, once for 10 min with wash buffer 4 (50 mM HEPES/KOH pH 7.5, 0.5 M NaCl, 0.1% sodium deoxycholate, 1% Triton X-100, 1 mM EDTA) at 4°C, and once for 10 min with wash buffer 5 (20 mM Tris-HCl pH 8, 250 mM LiCl, 0.5% NP-40, 0.5% sodium deoxycholate, 1 mM EDTA). Beads were then washed once with 1 mL resuspension buffer (50 mM Tris-HCl pH 7.5, 50 mM NaCl) and resuspended in 100 μL resuspension buffer before freezing at -80°C.

### LC/MS/MS Analysis

Samples were analyzed in a randomized order by LC-MS/MS using a Vanquish Neo coupled to an Orbitrap Astral mass spectrometer (ThermoFisher Scientific). A pooled sample was analyzed three times throughout the sample set. Samples were injected onto an IonOptics Aurora series 2 C18 column (75 μm id × 15 cm, 1.6 μm particle size) (IonOpticks) and separated over a 30 min method. The gradient for separation consisted of 2-30% mobile phase B at a 300 nL/min flow rate, where mobile phase A was 0.1% formic acid in water and mobile phase B consisted of 0.1% formic acid in 80% acetonitrile. Astral was operated in product ion scan mode for Data Independent Acquisition (DIA).

A full MS scan (m/z 380-980) was collected; resolution was set to 240,000 with a maximum injection time of 5 ms and AGC target of 500%. Following the full MS scan, a product ion scan was collected (30,000 resolution) and consisted of stepped higher collision dissociation (HCD) set to 25; AGC target set to 500%; maximum injection time set to 55 ms; variable precursor isolation windows 380-980 m/z; the isolation window was set to 3 m/z.

### TurboID-based Proteomics Data Analysis

Raw data files were processed using the directDIA method in Spectronaut (v20; Biognosys) and searched against the Uniprot reviewed *Saccharomyces cerevisiae* database (ATCC 204508/S288c, containing 6,727 entries, downloaded May 2024), the MaxQuant common contaminants database (246 entries), and yeast Tpk1/2/3 fused to TurboID. The following settings were used: enzyme specificity set to trypsin, up to two missed cleavages allowed, methionine oxidation, and N-terminal acetylation set as variable modifications. A false discovery rate (FDR) of 1% was used to filter all data. Cross-run normalization was enabled, imputation was disabled. Imputation and statistical analysis were performed in Perseus software (version 1.6.14.0). Proteins with log2 fold change ≥ 1 and a *p* < 0.05 are considered significant.

### Gene Ontology Analysis and Integration Analysis

Gene ontology (GO) analysis was conducted using g:Profiler. The gene symbols of protein interactors, except Tpk1/2/3, were queried with settings: *Organism = Saccharomyces cerevisiae*, *Statistical domain scope = Only annotated genes*, *Significance threshold = Benjamini-Hochberg FDR*, *Threshold = 0.05*. Integration analysis was performed using the module ‘joint pathway analysis’ of MetaboAnalyst (http://metaboanalyst.ca). DRMs from a single Tpk strain (e.g., Tpk1-only) were submitted as compounds, and interactors of other two Tpks (e.g., Tpk2/3) were submitted as proteins. Integration analysis was performed on ‘Metabolic pathways’ with settings: *Enrichment analysis = Hypergeometric Test*, *Topology measure = Degree Centrality*, *Integration method = Combined queries*.

### Data deposition

Mass spectrometry proteomics data have been deposited to the ProteomeXchange Consortium via the PRIDE [160] partner repository with the dataset identifier PXD067402.

Metabolomics data have been deposited to Metabolomics Workbench and can be accessed using the following account details: https://www.metabolomicsworkbench.org/data/DRCCMetadata.php?Mode=Study&DataMode=FactorsData&StudyID=ST001786&StudyType=MS&ResultType=5#DataTabs.

---

**Supplemental Figure 1. Paralogs have distinct roles in glucose sensing reflected by *pHXT5*-mNeonGreen. (A)** Left: promoter of *HXT5* was selected to drive expression of mNeonGreen. The reporter cassette was integrated at *YPRCtau3* locus in Chromosome XVI. Right: Induction of *pHXT5*-mNeonGreen in wildtype BY4741 over 4 h. Low glucose: SC medium with 0.05% glucose; High glucose: SC medium with 2% glucose. **(B)** Transcription reporter assays using *pHXT5*-mNeonGreen. Wildtype and mutant strains were incubated in high or low glucose for 4 h. The reporter intensities from mutant strains were normalized to that from wildtype and transformed by log_10_. Error bars show standard deviation for three independent experiments; each symbol represents the mean of four wells. Two-tailed Student t-tests were done to compare intensities between mutants and wildtype. *: *p* < 0.05; **: *p* < 0.01. **(C)** Time course data for one representative experiment for deletion of *RAS1/2*, *IRA1/2*, *GPB1/2*. Error bars show standard deviation for four wells representing the means of ∼1500 cells per well.

**Supplemental Figure 2. Inducible degradation of paralogs by IAA.** Top: Western blots validating expression of paralog-AID fusions and degradation of paralogs upon IAA addition. Cells were treated with 150 μM IAA or same volume of DMSO for 30 min before harvest. Bottom: Ponceau stain.

## ACKNOWLEDGEMENTS

Funded by NIH grant R35-GM118105 (H.G.D.) and R35GM119518 (D.I.). We thank Matthew P. Miller at University of Utah for kindly providing plasmids and Joseph C. Reese at The Pennsylvania State University for providing protocols for the BioID experiments. We thank Angie L. Mordant for mass spectrometry data analysis of BioID experiments as well as Dr. Yuan Li and support staff in the Metabolomics and Exposome Laboratory for their assistance. This research is based in part upon work conducted using the UNC Proteomics Core Facility, which is supported in part by NCI Center Core Support Grant (2P30CA016086-45) to the UNC Lineberger Comprehensive Cancer Center. The UNC Metabolomics and Exposome Laboratory at the UNC Nutrition Research Institute conducted the untargeted metabolomics analysis under funding from the Sumner laboratory.

Y.S. and H.G.D. were responsible for conceptualization. Y.S. and H.G.D. designed experiments. Y.S., K.A., and D.I. constructed strains and plasmids. Y.S., K.A. performed transcription reporter assays and data analysis. D.I. performed pH recovery experiments and data analysis. S.L.M. and B.R.R. performed sample preparation and data collection for metabolomics. S.J.S., S.L.M. and B.R.R. performed data analysis for metabolomics. Y.S., S.P.L. and performed sample preparation, data collection and data analysis of BioID-proteomic study. Y.S. and H.G.D. were responsible for writing, reviewing and editing the manuscript.

